# A Cytoskeletal Function for PBRM1 Reading Methylated Microtubules

**DOI:** 10.1101/2020.04.21.053942

**Authors:** Menuka Karki, Rahul K. Jangid, Riyad N.H. Seervai, Jean-Philippe Bertocchio, Takashi Hotta, Pavlos Msaouel, Sung Yun Jung, Sandra L. Grimm, Cristian Coarfa, Bernard E. Weissman, Ryoma Ohi, Kristen J. Verhey, H. Courtney Hodges, Ruhee Dere, In-Young Park, W. Kimryn Rathmell, Cheryl L. Walker, Durga N. Tripathi

## Abstract

The chromatin modifier SETD2 was recently shown to be a dual-function methyltransferase that “writes” methyl marks on both chromatin and the mitotic spindle, revealing α-tubulin methylation as a new posttranslational modification of microtubules. Here, we report the first cytoskeletal “reader” for this SETD2 methyl mark: the polybromo protein PBRM1. We found PBRM1 directly binds the α-Tub-K40me3 mark on tubulin, and localizes to the mitotic spindle and spindle pole during cell division. PBRM1 can assemble a PBAF complex in the absence of chromatin as revealed by mass spectrometry, and can recruit other PBAF complex components including SMARCA4 and ARID2 to α-tubulin. In addition to PBRM1, other PBAF components were also localized to the mitotic spindle and spindle pole. This PBAF localization was dependent on recruitment to microtubules by PBRM1, and loss of spindle-associated PBRM1/PBAF led to genomic instability as assessed by increased formation of micronuclei. These data reveal a previously unknown function for PBRM1 beyond its role remodeling chromatin, and expand the repertoire of chromatin remodelers involved in writing and reading methyl marks on the cytoskeleton. The results of this study lay the foundation for a new paradigm for the epigenetic machinery as chromatocytoskeletal modifiers, with coordinated nuclear and cytoskeletal functions.

## Introduction

PBRM1 (Polybromo-1), also known as PB1 or BAF180, is a subunit of the PBAF (Polybromo BRG1 Associated factor) chromatin remodeler complex that belongs to the SWI/SNF family^1^ in mammalian cells, with homology to RSC complex in yeast^2^. PBRM1 along with two other subunits, BRD7 and ARID2, distinguish PBAF from other SWI/SNF complexes, termed BAF and ncBAF^3^. PBAF and other SWI/SNF ATP-dependent chromatin remodeling complexes have emerged as important drivers of cancer, with genes encoding SWI/SNF subunits mutated in approximately 20% of all cancers^4,5^. *PBRM1* mutations are particularly frequent in clear cell renal cell carcinoma (ccRCC), with one allele of *PBRM1* lost in >95% of ccRCC, and defects in the second *PBRM1* allele occurring in >40% cases of ccRCC, making *PBRM1* the second most common oncogenic driver of ccRCC after *VHL* (Von Hippel-Lindau)^6,7^.

Recently, we reported that in addition to histones, the methyltransferase *SETD2* (third most frequent driver for ccRCC) also methylates lysine 40 of alpha tubulin (α-TubK40me3) ^8,9^. The α-TubK40me3 mark localizes to spindle microtubules during mitosis and to the midbody during cytokinesis. Loss of SETD2 and α-TubK40me3 causes genomic instability and defects such as multipolar spindle formation, chromosomal bridges at cytokinesis, micronuclei, polyploidy, and polynucleation; a phenotype specifically-linked SETD2 activity as a tubulin methyltransferase ^8,9^. However, how the α-TubK40me3 mark on microtubules is “read” to maintain genomic stability is not known. Here, we report PBRM1 reads the SETD2-derived α-TubK40me3 mark on microtubules during mitosis, recruiting other PBAF complex components including SMARCA4 and ARID2 to the mitotic spindle to maintain genomic stability.

## Results

### PBRM1 binds α-tubulin and localizes to spindle microtubules during mitosis

We found while PBRM1 localized primarily to the nucleus during interphase, in mitotic cells PBRM1 localization was highly enriched at the poles of the mitotic spindle. This localization pattern was observed using either a GFP-tagged PBRM1 fusion protein or an antibody directed against endogenous PBRM1 in several cell types including human kidney epithelial (HKC), 786-0 (ccRCC) and HEK293T cells (**Fig. 1a-c**). Immunoreactivity at the spindle poles was specific for PBRM1 as this pattern of localization was seen in PBRM1-proficient but not PBRM1-deficient cells (**Fig. 1b**). To further confirm spindle localization of PBRM1 during mitosis, we performed live-cell imaging of GFP-PBRM1 in HEK293T cells, which clearly showed PBRM1 localizing to the mitotic spindle, with most intense localization at the spindle pole (**Supplementary Video 1**).

**Fig. 1:**
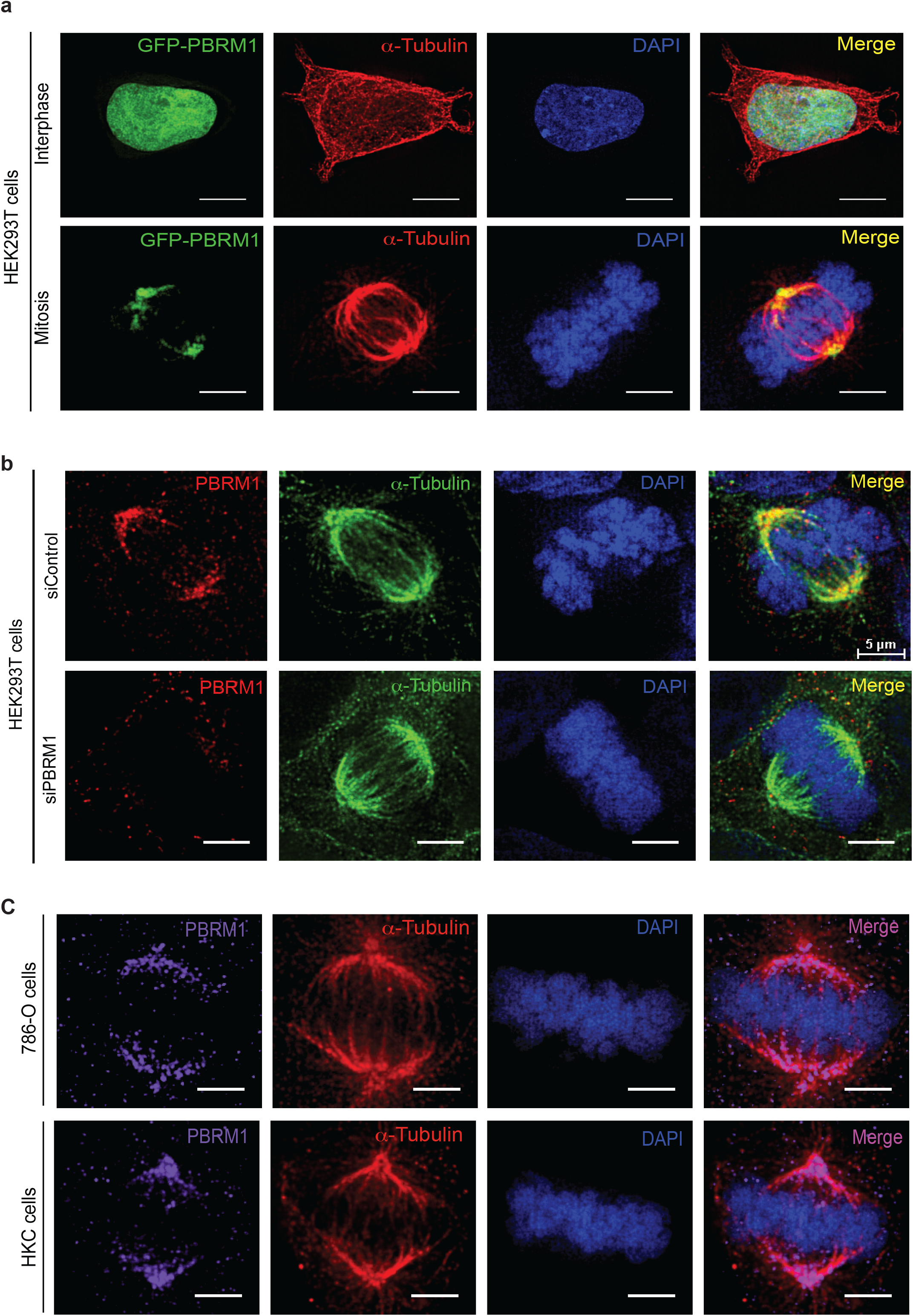
PBRM1 localizes with spindle microtubules during mitosis: **a**, representative images from deconvolution microscopy of HEK293T cells (during interphase and mitosis) ectopically expressing GFP-PBRM1 (green), stained with an antibody to α-tubulin (red) and DAPI to visualize chromosomes (blue). Scale bars, 10μm (upper panel) and 5μm (lower panel). (n=3). **b**, Representative deconvolution images of HEK293T cells stained with antibodies for endogenous PBRM1 (red), α-tubulin (green) and DAPI to visualize chromosomes (blue) showing subcellular localization of PBRM1 at the mitotic spindle (upper panels) lost in PBRM1 depleted HEK293T cells (lower panel). Scale bars 5μm. (n=3). **c**, Representative images from deconvolution microscopy showing localization of PBRM1 to the mitotic spindle in 786-O and HKC cells stained for α-tubulin(red) and DAPI to visualize chromosomes (blue) during mitosis. Scale bars, 5μm. (n=3 biological replicates for both HKC and 786-O cells).

Next, we performed microtubule co-sedimentation assays using lysates prepared under low-stringency conditions from parental and *PBRM1* CRISPR-KO HEK293T cells, which showed PBRM1 co-sedimented with taxol-stabilized microtubules in the pelleted fraction, thus confirming the specificity of the PBRM1 association with microtubules (**Fig. 2a**). Finally, we performed co-immunoprecipitation experiments in HEK293T, 786-O and MEFs cells to demonstrate PBRM1 and α-tubulin could be co-immunoprecipitated from cells using both GFP-tagged and endogenous PBRM1 (**Fig. 2b-e)**. These data revealed PBRM1 is a tubulin-binding protein that localizes to spindle microtubules during mitosis.

**Fig. 2:**
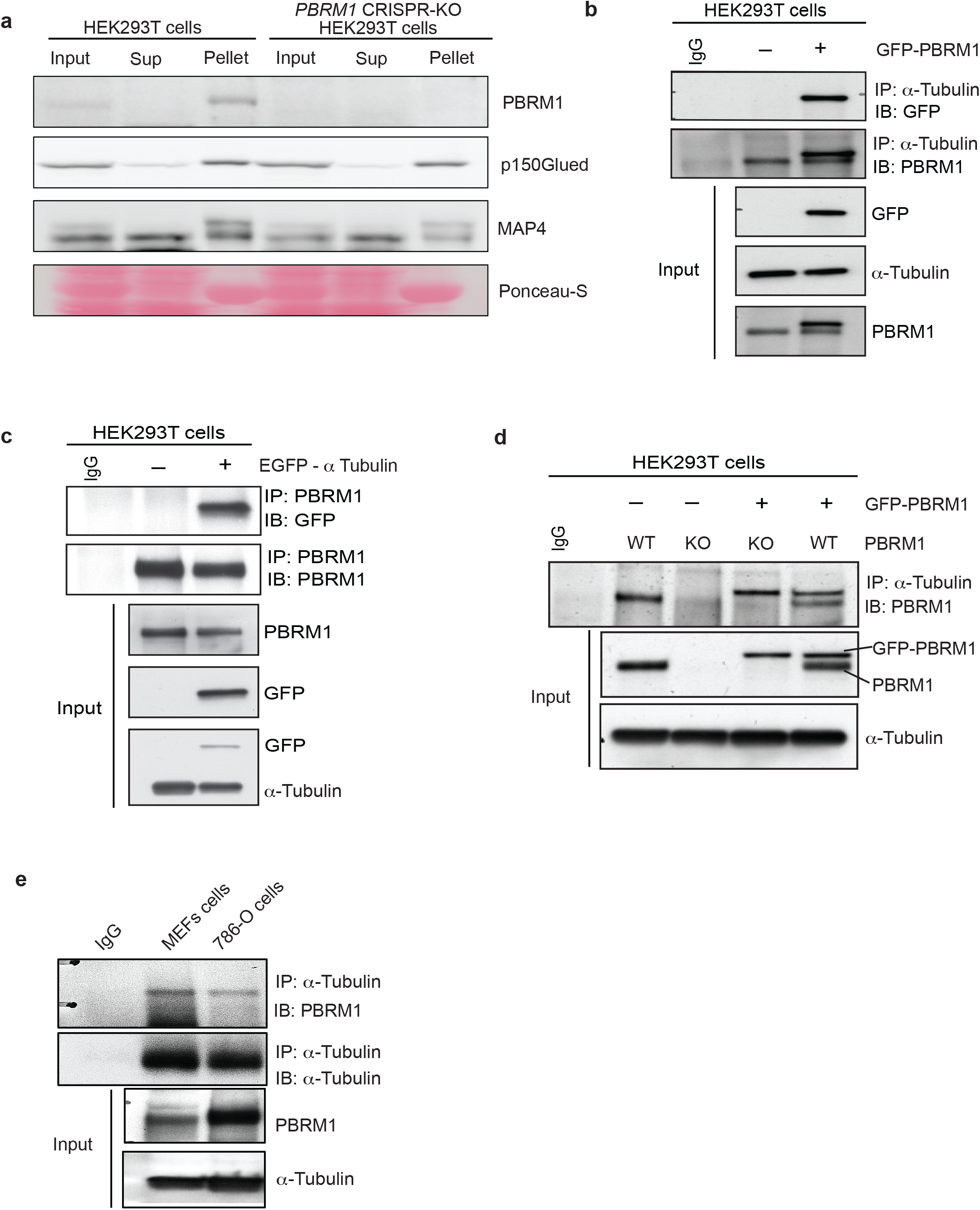
PBRM1 interacts with α-tubulin: **a,** Western analysis following microtubule co-sedimentation assay of parental HEK293T or *PBRM1* CRISPR-KO HEK293T cells using a PBRM1 antibody; MAP4 and p150Glued antibodies used as positive control known microtubule associated proteins **b**, Immunoblot analysis showing co-immunoprecipitation (Co-IP) of endogenous α-tubulin and ectopically expressed PBRM1 and respective input lysates from HEK293T cells. **c**, Immunoblot analysis showing Co-IP of endogenous PBRM1 and ectopically expressed α-tubulin and respective input lysates from HEK293T cells. Representative blots (n=3). **d**, Immunoblot analysis in HEK293T cells for co-immunoprecipitation of PBRM1 and α-tubulin and their corresponding input lysates from PBRM1 proficient-, PBRM1-depleted, and PBRM1-depleted cells rescued with PBRM1 showing specificity for tubulin interaction with PBRM1. Representative blots (n=3). **e** Immunoblot analysis of co-immunoprecipitation of endogenous α-tubulin and PBRM1 and corresponding input lysates from mouse embryonic fibroblasts (MEFs) and 786-O cells. Representative blots (n=3).

### PBRM1 recognizes methylated α-tubulin

We found PBRM1 co-localized with the α-TubK40me3 mark on spindle microtubules made by the SETD2 methyltransferase^8,9^ as seen by co-localization of an antibody directed against the α-TubK40me3 mark and GFP-tagged PBRM1 (**Fig. 3a**) and confirmed by overlap of the peak intensity of line profiles through the mitotic spindle (**Fig. 3b**). This immunocytochemistry data suggested PBRM1 binds to methylated α-tubulin and to test this, we expressed EGFP-tagged wild-type, K40Q (acetylation mimic), and K40R (methyl- and acetyl-deficient) mutant α-tubulin in HEK293T cells and found PBRM1 association with α-tubulin was dramatically reduced by both the K40R and K40Q mutations (**Fig. 3c**).

**Fig. 3:**
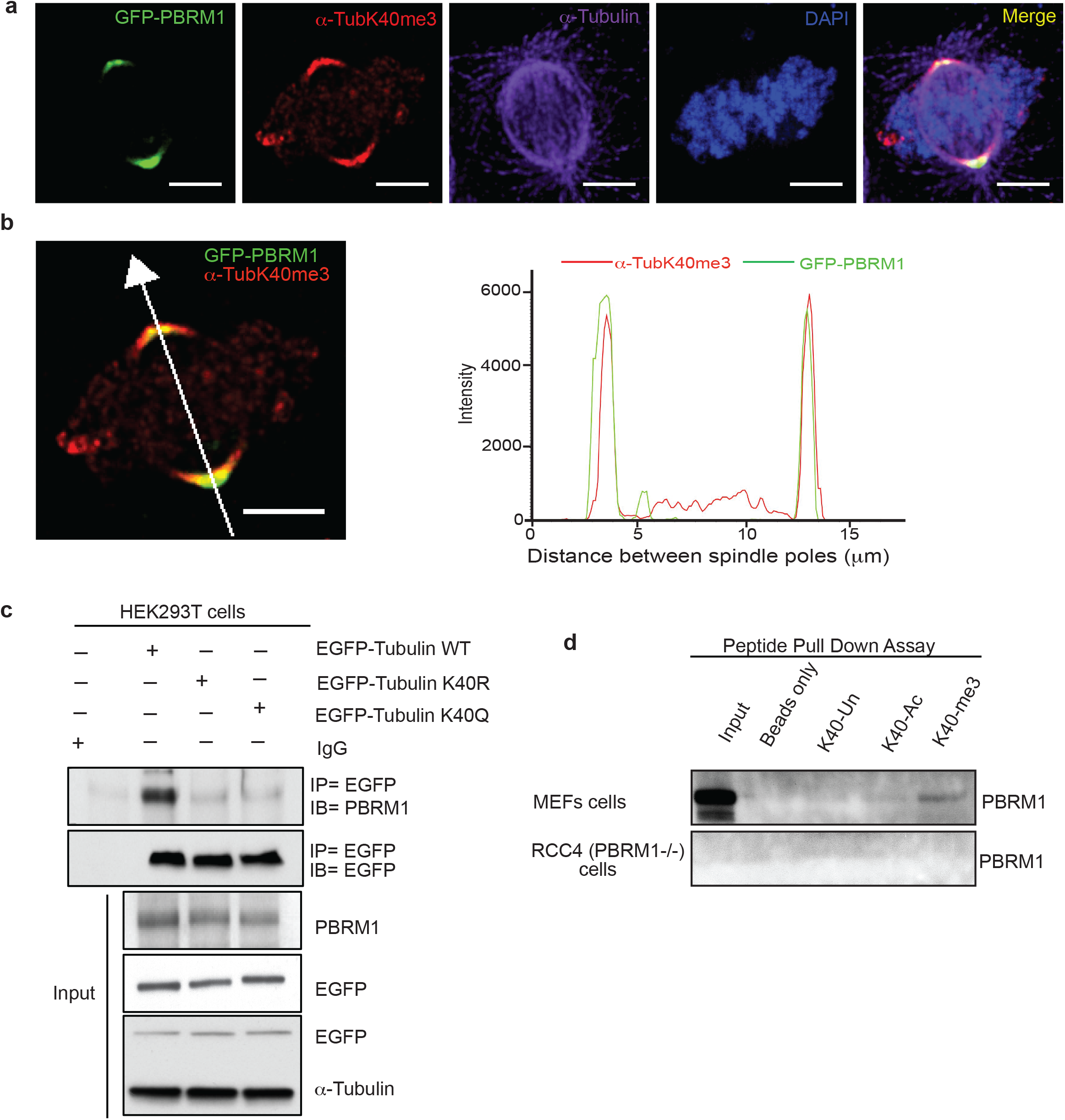
PBRM1 localizes with methylated α-tubulin during mitosis: **a**, Representative images of HEK293T cells ectopically expressing GFP-PBRM1 (green) and stained with antibodies specific for the SETD2 methyl mark (α-TubK40me3 red), α-tubulin (far red/purple) and DAPI to visualize chromosomes (blue) showing PBRM1 co-localization with α-TubK40me3 and α-tubulin at mitotic spindle and spindle pole. **b,** Colocalization of PBRM1 (green) and α-TubK40me3 (red), line profiles were obtained between the two spindle poles using deconvolution microscope NIS elements software, which showed green and red signals are aligned at spindle pole and PBRM1 and a-Tub-K40me3 are co-localized. Scale bar 5μm. (n=3 biological replicates). **c,** Immunoblot analysis following co-immunoprecipitation of PBRM1 and ectopically expressed WT or mutant α-tubulin (K40R) or acetylation mimic (K40Q) and respective input lysates from HEK293T cells. Representative blots (n=3). **d**, Immunoblot analysis following peptide pull down using biotin labelled K40 α-tubulin peptides that were unmodified (K40-UN), acetylated (K40-Ac) or tri-methylated (K40-me3) using lysates from PBRM1 proficient (MEF) or deficient (RCC4) cells. Representative blots shown (n=3).

To confirm PBRM1 binding to α-tubulin was α-TubK40me3 dependent, we synthesized α-tubulin peptides with unmodified, acetylated, or trimethylated lysine 40, and performed peptide pull down assays, which confirmed PBRM1 bound to methylated, but not acetylated or unmodified, α-tubulin K40 peptides (**Fig. 3d**). Finally, to confirm binding of PBRM1 to microtubules occurred via recognition of the SETD2 α-TubK40me3 mark, using SETD+/+ and SETD2−/− 786-O cells we found co-localization of PBRM1 at the spindle poles was dramatically reduced in *SETD2*-null cells (**Fig. 4a**). This reduction of PBRM1 binding was confirmed by line profiles through the mitotic spindle to quantitate the peak intensity of the PBRM1 signal (**Fig. 4b**), which showed that the most intense focal binding of PBRM1 at the spindle poles was significantly reduced in SETD2 null cells (**Fig. 4c).** Taken together, these data identify PBRM1 as a microtubule binding protein that recognizes the SETD2 α-TubK40me3 mark during mitosis.

**Fig. 4.**
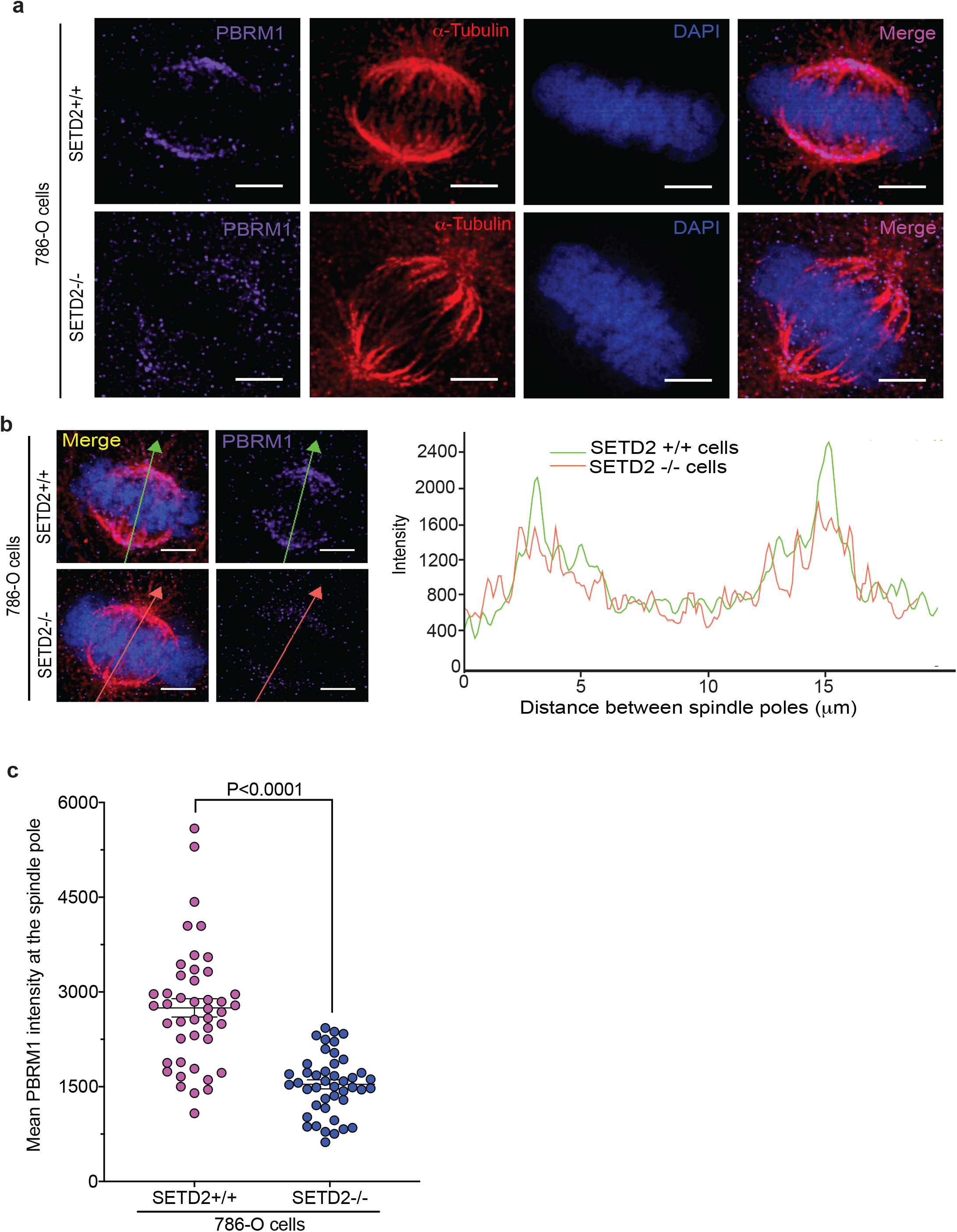
Dependency on the SETD2 α-TubK40me3 mark on microtubules for PBRM1 interaction with α-tubulin: **a,** Representative images from deconvolution microscopy showing localization of PBRM1 to the mitotic spindle in *SETD2*+/+ and *SETD2*−/− 786-O cells during mitosis stained for PBRM1 (purple) and α-tubulin (red) and DAPI to visualize chromosomes (blue). Scale bars 5μm. (n=3 biological replicates). **b**, Representative images of line intensity profiles and intensity measurement of PBRM1 on mitotic spindle in *SETD2*+/+ *vs SETD2*−/− 786-O cells using deconvolution microscope NIS elements software. Scale bars, 5μm. (n=3). **c**, Quantification of intensity at peak spindle pole localization of PBRM1 from **b** in *SETD2*+/+ *vs SETD2*−/− 786-O cells. Data is represented as mean ± S.E.M. p-value determined by t-test (n= 45 mitotic cells/condition) followed by Mann Whitney posthoc test.

### PBRM1 recruits SWI/SNF (PBAF) subunits to methylated microtubules

SWI/SNF components typically co-exist as a complex in cells, and the presence of one suggests the presence of others^10^. We thus asked whether PBRM1 might be recruiting other components of the PBAF complex to mitotic microtubules. Both PBRM1 and the ATPase subunit of PBAF, SMARCA4/BRG1, could be detected in the cytoplasmic fraction of cells by western analysis (**Fig 5a**), prompting us to perform mass spectrometry from this subcellular compartment to identify PBAF components that could associate with PBRM1 in the absence of chromatin. Following immunoprecipitation of PBRM1 from the cytoplasmic fraction of HEK293T cells, we confirmed by mass spectrometry PBRM1 exists in a chromatin-independent complex with SMARCA4, ARID2, BRD7, SMARCB1, SMARCC1 and other PBAF components (**Fig 5b,c and Table 1)**. Consistent with PBRM1 binding microtubules as part of a PBAF complex, SMARCA4 and ARID2 could also be co-immunoprecipitated with α-tubulin (**Fig. 5d**), and both localized to the spindle pole during mitosis (**Fig. 5e upper panel**); knockdown of PBRM1 abrogated spindle localization of both SMARCA4 and ARID2 (**Fig. 5e lower panel**).

**Table 1:** iBAQ3 values obtained by mass spectrometry from immunoprecipitated samples using PBRM1 antibody from HEK293T cells and *PBRM1* CRISPR-KO HEK293 cells cytoplasmic fraction.

| Groups | Average iBAQ |  |  |  |
| --- | --- | --- | --- | --- |
|  | Cytoplasmic fraction of HEK293T cells |  | Cytoplasmic fraction of PBRM1 CRISPR-KO HEK293T cells |  |
|  | Pre-Clear | PBRM1-IP | Pre-Clear | PBRM1-IP |
| PBRM1 (BAF180) | 0.3 | **462.0** | n.d. | 0.7 |
| ARID2 | n.d. | **219.0** | 0.2 | 0.3 |
| BCL7A | n.d. | **52.9** | n.d. | 0.0 |
| BCL7C | n.d. | **48.6** | n.d. | 0.0 |
| BRD7 | n.d. | **80.1** | 0.8 | 0.0 |
| PHF10 | n.d. | **53.3** | n.d. | 0.0 |
| SMARCA4 (BRG1) | n.d. | **173.0** | 1.9 | 0.1 |
| SMARCB1 (BAF47) | 4.3 | **125.5** | 7.7 | 0.9 |
| SMARCC1 (BAF155) | n.d. | **120.0** | 2.5 | 1.1 |
| SMARCC2 (BAF170) | 1.2 | **122.5** | 5.5 | 0.6 |
| SMARCD1 (BAF60A) | 0.4 | **43.3** | 0.8 | 0.0 |
| SMARCD2 (BAF60B) | 0.4 | **114.5** | 4.6 | 0.0 |

**Fig. 5.**
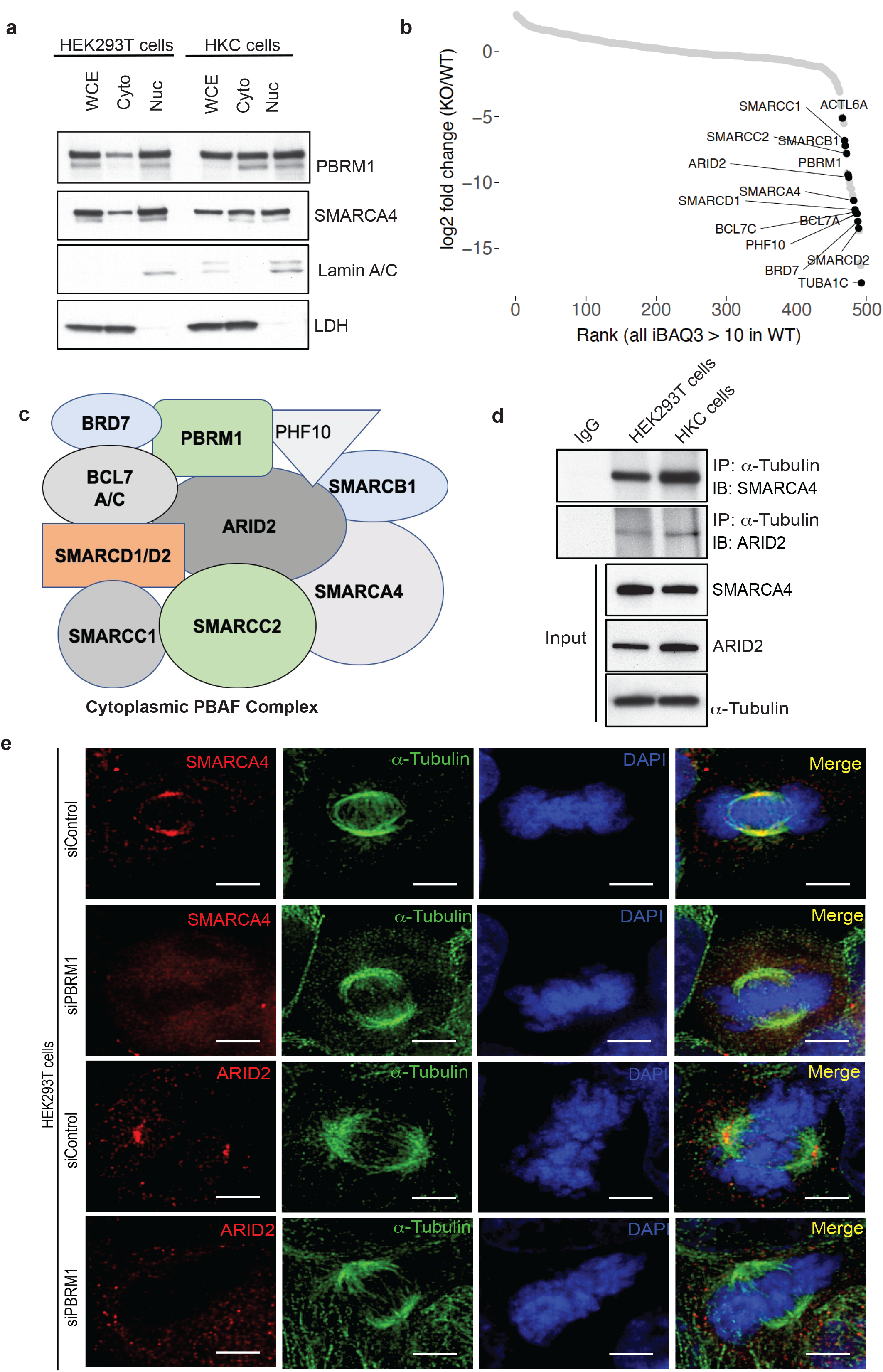
PBRM1 complex formation with SWI/SNF (PBAF) subunits is chromatin-independent and recruits SWI/SNF (PBAF) subunits to methylated microtubules: **a**, Immunoblot analysis showing PBRM1 and SMARCA4 in the cytoplasmic as well as nuclear fraction of HEK293T and HKC cells. LDH and Lamin A/C were used as markers for cytoplasmic (Cyto) and nuclear (Nuc) fractions respectively. Representative blots (n=3). **b**, Quantification of PBAF-specific subunits complexed with PBRM1 in PBRM1-proficient cells after PBRM1 immunoprecipitation from the cytoplasmic fraction of cell lysates from HEK293T cells vs *PBRM1* CRISPR-KO HEK293T cells followed by mass spectrometry. **c.** Schematic of SWI/SNF subunits complexed with PBRM1 identified by mass spectrometry using cytoplasmic fraction of HEK293T cells from (b). **d.** Immunoblot analysis showing co-immunoprecipitation of SMARCA4 and ARID2 with α-tubulin and their corresponding Input lysates from HEK293T and HKC cells. Representative blot (n=3). **e,** Representative image using deconvolution microscopy of cells stained with antibodies directed against SMARCA4 (red, upper two panels) or ARID2 (red lower two panel) and α-tubulin (green) DAPI to visualize chromosomes (blue). Representative images show SMARCA4 and ARID2 localization at the mitotic spindle and spindle pole in PBRM1 proficient HEK293T cells (upper panel) and loss of mitotic spindle localization in PBRM1 depleted HEK293T cells (lower panel). Scale bars, 5μm. (n=3 biological replicates).

### Loss of PBRM1 disrupts genomic stability

In yeast, the PBRM1 homolog RSC2 is essential for chromosome arm cohesion^11,12^. Xue et al had localized PBRM1 to the kinetochore in the absence of microtubules^13^, and in a later study, the induction of genomic instability observed upon loss of PBRM1^14^ was thought to be due to loss of centromere cohesion. However, we did not see prominent localization of PBRM1 or SMARCA4 at the centromere of mitotic chromosomes with our staining protocol (**Fig. 1, 3, 4 and 5**), suggesting rather PBRM1 localization to the spindle pole was required for maintenance of genomic stability. To explore this possibility, we knocked down PBRM1 in HEK293T cells, which significantly increased formation of micronuclei; re-expression of PBRM1 rescued this genomic instability (**Fig 6a and b**). Interestingly, this phenocopied the genomic instability observed in *SETD2*-null cells that had lost the α-TubK40me3 mark^8,9^.

**Fig. 6:**
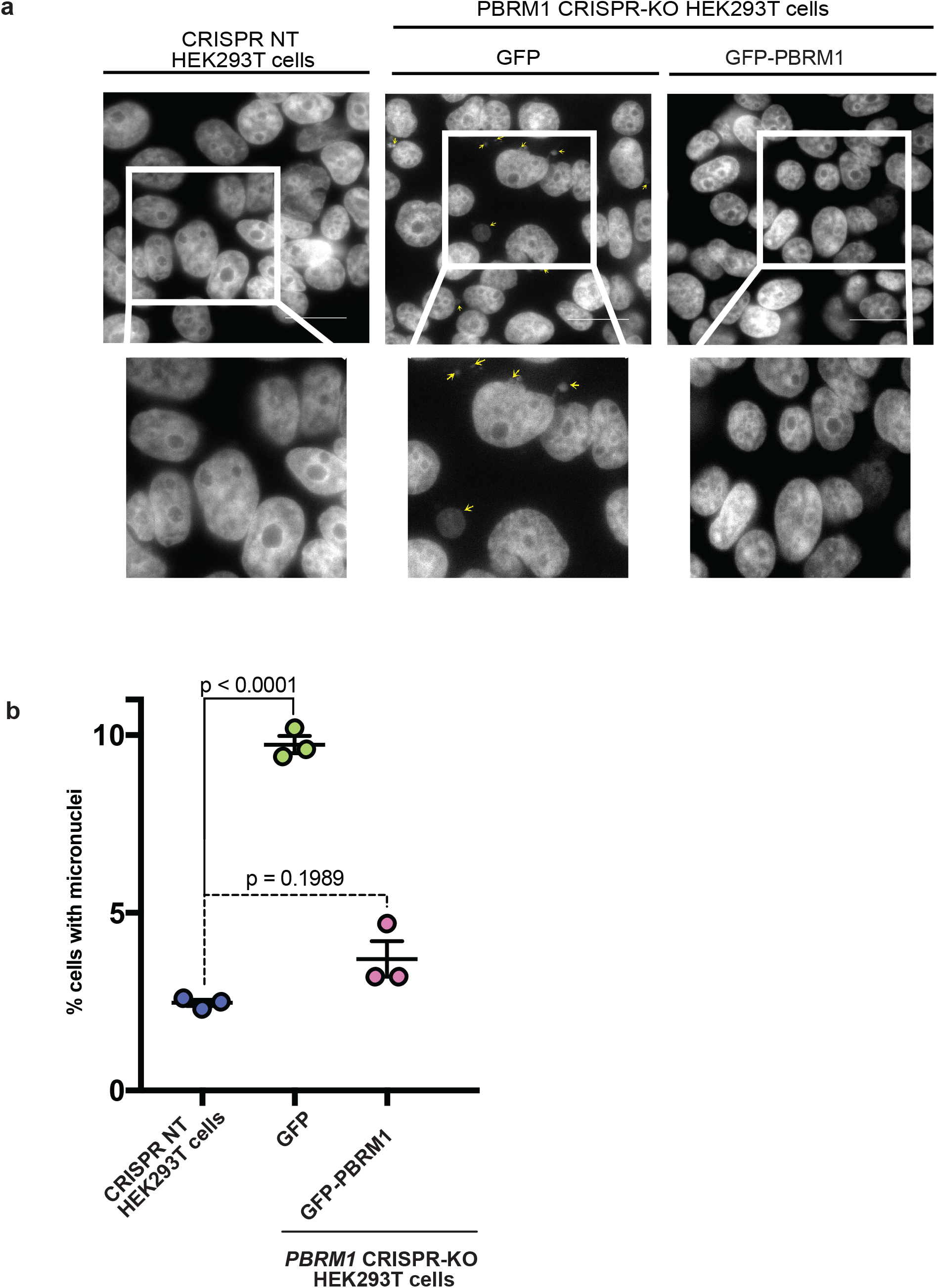
PBRM1 absence from mitotic spindle causes genomic instability: **a,** Representative images of micronuclei (indicated by small arrows) genomic instability phenotype observed with loss of PBRM1 or re-expression of α-tubulin binding-deficient PBRM1 versus expression of WT or GFP-PBRM1 α-tubulin binding-proficient PBRM1 in *PBRM1* CRISPR-KO HEK293T cells. Scale bars, 25μm. (n=3). **b**, Quantification of frequency of micronuclei containing cells from (a). Data are represented as mean ± S.E.M. for three independent biological replicates, with 1000 cells counted per replicate. p-value determined by one-way ANOVA followed by Dunnett’s posthoc test for multiple comparisons against control cells.

## Discussion and Conclusions

Recognition of the SETD2 α-TubK40me3 mark by PBRM1 provides a new framework for understanding the role of SETD2 and PBRM1 in maintaining genomic stability, and reveals the first functional linkage at the cytoskeleton for these two chromatin modifiers. Although, microtubules can be acetylated or methylated at lysine 40, our data show PBRM1 binds specifically to methylated α-tubulin, recruiting other component of PBAF complex and regulate genomic stability. These data also suggest a model whereby PBAF has dual chromatin and cytoskeleton activity via binding to PBRM1 (**Fig. 7**).

**Fig. 7:**
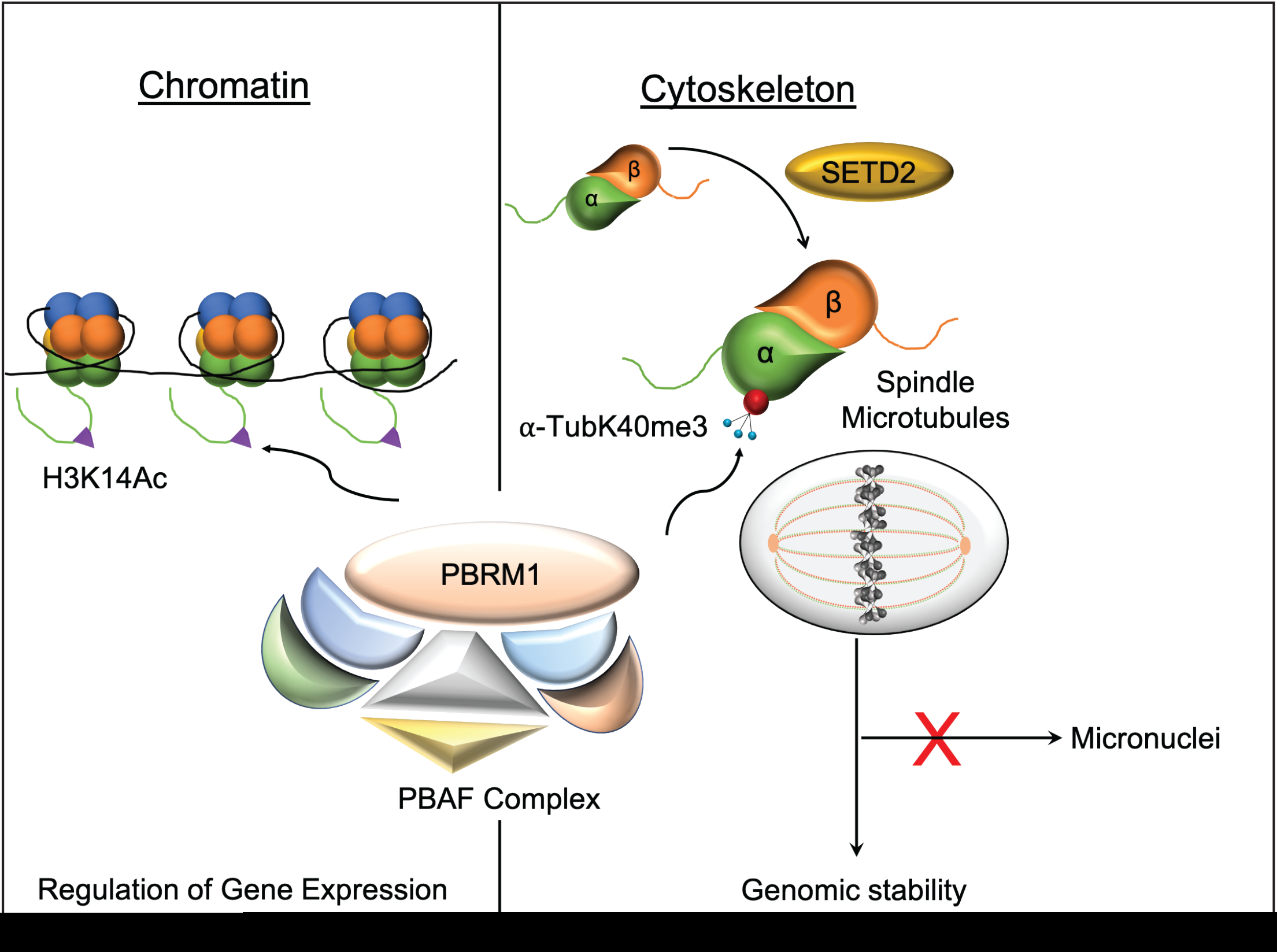
Graphical model showing dual chromatin and cytoskeletal function of PBRM1 in regulating gene expression and genomic stability via interactions with chromatin in the nucleus and microtubules of the mitotic spindle binding to α-TubK40me3, respectively.

An intriguing aspect of microtubule biology is the luminal positioning of lysine 40, the target for the α-TubK40me3 methyl mark made by SETD2 and the acetyl mark made by alpha tubulin acetyl transferase (ATAT-1). Luminal positioning of acetyl and methyl marks raises the question as to how writers, such as ATAT-1 and SETD2 access these residues, and how luminal acetyl and methyl marks are read to direct the structure and function of microtubules. Whether SETD2 methylates α-tubulin prior to or after assembly of α/β tubulin dimers into microtubules is not known. However, Even et al., reported that ATAT-1 is transported by vesicles along the length of microtubules, accessing K40 to acetylate this residue via pores in the microtubule shaft^15^. These and other studies have led to the growing appreciation that lattice dynamics of even stable microtubules can provide access to luminal residues along the shaft of microtubule polymers. Recently, Schaedel et al., reported a passive breathing mechanism for microtubules in which α/β tubulin dimers can be evicted and replaced from the microtubule shaft via an energy consuming process^16^. SWI/SNF complexes are known to evict histones and other proteins from nucleosomes during chromatin remodeling^17–19^, suggesting the intriguing hypothesis that PBAF, and perhaps other SWI/SNF complexes, could be playing a similar role in evicting tubulin and/or other microtubule associated proteins during microtubule remodeling.

Finally, co-localized on chromosome 3p, one allele each of *PBRM1* and *SETD2* are co-deleted as an early event in development of ccRCC^20^, with “second hits” in these tumor suppressors the second and third most common oncogenic events in this cancer. In a recent evolutionary model of ccRCC developed by the TRACERx study^21,22^, second hits resulting in loss of the ‘reader’ PBRM1 precede loss of the ‘writer’ SETD2; in no case has loss of SETD2 (and its methyl mark) preceded PBRM1 loss. While the reason for this sequence has remained obscure, our results suggest that cancer cells with second hits that inactive SETD2 will lack the cytoskeletal methyl-mark read by PBRM1, thereby abrogating PBRM1 cytoskeletal function, and may gain no further (cytoskeletal) advantage by loss of PBRM1. Conversely, while early loss of PBRM1 can contribute to tumor progression by eliminating an α-TubK40me3 reader, later loss of SETD2 will result in loss of both the cytoskeletal α-TubK40me3 and chromatin H3K36me3 marks, abrogating SETD2 chromatin remodeling ability, and potentially, the activity of any other ‘readers’ of its cytoskeletal methyl mark. Of note, in the setting of loss of PBRM1, repeated subclonal selection for SETD2 mutations occurs^22^, suggesting that complete loss of both cytoskeletal and chromatin methyl-marks may be an evolutionary bottleneck in cancer progression. Thus, our study identifies dysregulation of microtubule methylation as novel nexus of convergence in explaining chromatocytoskeletal “writer” (SETD2) and “reader” (PBRM1) can drive genomic instability, opening new windows into understanding mechanisms by which chromatin remodeler defects can drive development of cancer and other diseases.

## Acknowledgements

We would like to acknowledge Dr. Hugo Bellen’s lab at Baylor college of medicine for kindly providing the H2B-mRuby2 plasmid. This work is supported by grants from the Department of Defense: KC170259 (D.N.T.); the Owen’s Foundation (D.N.T.); the National Institutes of Health: NCI-R35CA231993 (C.L.W.) and R01CA203012 (W.K.R., C.L.W.), the Templeton Foundation: #61099 (C.L.W.). R.N.H.S. is supported by the American Heart Association (19PRE34430069), the Baylor College of Medicine (BCM) Medical Scientist Training Program, and a BP America Biomedical Scholarship from the BCM Graduate School of Biomedical Sciences. J.P.B. was funded by La Ligue Nationale Contre le Cancer and the Philip Foundation. P.M. is supported by a Young Investigator Award from the Kidney Cancer Association, a Career Development Award by the American Society of Clinical Oncology / Conquer Cancer Foundation, a Concept Award by the Department of Defense, and a Khalifa Scholar Award by the Institute for Personalized Cancer Therapy (The University of Texas MD Anderson Cancer Center). National Institutes of Health: P30CA125123, NCI center grant and Cancer Prevention and Research Institute of Texas (CPRIT): RP170005 for Mass Spectrometry Proteomics Core. KJV is supported by NIH R35GM131744.

## Contributions

D.N.T., I.Y.P. and C.L.W. conceptualized the study. D.N.T., C.L.W., I.Y.P., R.D. and P.M. designed experiments, with input from H.C.H., K.J.V., W.K.R. and R.O. Further, D.N.T., R.N.H.S., M.K., R.J., J.P.B., H.C.H, S.L.G., C.C., and C.L.W. wrote the manuscript with input from all authors. D.N.T. performed experiments, with assistance as follows: M.K. performed immunofluorescence experiments. R.J. performed biochemical experiments. H.C.H. assisted for image analysis and quantification. T.H. performed microtubule co-sedimentation experiments, with supervision from R.O. S.Y.J. assisted with mass spectrometry experiments. Unless noted, all work was performed under supervision of D.N.T. and C.L.W.

## Data and Material availability

All data are available in the main text and supplementary materials.

## Materials and Methods

### Antibodies and Reagents

The details of the antibodies used for Immunoblotting (IB) and Immunofluorescence (IF) are given below PBRM1 (Bethyl Lab, # A301-591A, 1:1,000 IB), Lactate Dehydrogenase (LDH) (Abcam, # ab47010, 1:5,000 IB), Lamin A/C (Cell signaling, # 2032S, 1:1,000 IB), DM1A (Santa Cruz, cat# sc32293, 1:4,000 IB, IF), GFP (Santa Cruz, # sc-9996, 1:5,000 IB), p150Glued (BD Biosciences, 610474, 1:400 IB), MAP4 (Abcam, # ab89650, 1:800 IB), α-TubK40me3 (Custom made Rabbit polyclonal antibody from Covance, 1:1000 IB), SMARCA4 (Santa Cruz, # sc-17796 IB), ARID2 (Santa Cruz, # sc-166117,1:500 IB), PBRM1 (Bosterbio, # M01130, 1:200 IF), SMARCA4 (ThermoFisher Scientific, # 720129, 1:200 IF), α-tubulin (Sigma-Aldrich, # T6199, 1:4,000 IB), α-tubulin (ThermoFisher Scientific, # PA5-19489, 1:5,000 IF) and ARID2 (Novus biologicals, # NBP-2-57220, 1:250 IF). Secondary antibodies (1:5,000) conjugated to horseradish peroxidase were purchased from Santa Cruz Biotechnology. For immunofluorescence staining, Alexa Fluor-labelled secondary antibodies (1:2,000) were purchased from Invitrogen. Nuclear counterstaining reagent, DAPI (1:4,000), was purchased from Invitrogen. SlowFade Gold Antifade reagent (ThermoFisher Scientific, # S36937) was used as mounting medium for coverslips.

### Cell Culture and transfection

Human Embryonic Kidney (HEK) 293T cell lines acquired from ThermoFisher Scientific (# R70007) and Human Kidney Cells (HKC) (acquired from Collaborator Dr. Kim Rathmell Lab, Vanderbilt University, TN) were maintained in DMEM (Sigma-Aldrich, # D6429) media supplemented with 10% FBS. The human 786-0 *SETD2+/+ and SETD2 −/−* RCC cell line (acquired from Collaborator Dr. Kim Rathmell Lab, Vanderbilt University, TN) were maintained in DMEM medium (Life Technologies, #11875-085) supplemented with 10% FBS. Mouse Embryonic Fibroblast (MEFs) cells were maintained in Phenol free DMEM media (Gibco, # 21063-029) supplemented with 10% FBS, Sodium pyruvate (Gibco, #11360070) and Glutamax (Gibco, # 35050061). All the FBS used to supplement the media were purchased from Sigma (# F24429). The transient transfection of plasmids and siRNAs were performed using Lipofectamine 2000 reagent (ThermoFisher Scientific, # 11668-500) and Dharmafect-1 (Dharmacon, # T-2001-03) respectively, according to the manufacturer’s instructions. All of the cell lines used in this study were routinely tested and confirmed negative for mycoplasma.

### Plasmid DNA preparation

GFP-PBRM1 (Addgene, # 65387) plasmids were gifts from Kyle Miller ^23^. pEGFP-Tubulin K40R (Addgene plasmid # 105303) and K40Q (Addgene plasmid # 105302) plasmids were gifts from Kenneth Yamada^24^. Plasmids from Addgene were grown from single clones which were sequence verified, and maxipreps ((Omega **Bio-tek, inc. D6922-04)** to generate plasmid DNA for transfection.

### siRNA knockdown of PBRM1

ON-TARGET plus SMART pool Human-PBRM1 siRNA (# L-008692-01-5) and ON-TARGET plus Non-Targeting pool (# D-001810-10-20) were purchased from Dharmacon. siRNAs were resuspended in 1X siRNA buffer (GE Dharmacon) to obtain 20 μM stock. HEK293T cells were transfected with indicated siRNA at 10nM final concentration with Dharmafect-1 according to the manufacturer’s instructions for 72-hour prior lysates collection or prior fixing cells for immunostaining.

### Generation of PBRM1-KO HEK293T stable cell lines using CRISPR-Cas9 and validation

PBRM1 KO HEK293T cells were generated using guide RNA with CAS9 construct purchased from Genecript. Guide RNA sequence (GAAACCACTTCATAATAGTC) was designed on exon 4. Cells were transfected with CAS9 construct for 48 hours and put on puromycin selection (2μg/ml) for one week to isolate immortalized Non target (NT) and/or PBRM1-KO HEK293T cells. Isolated colonies were picked using cloning rings after one week of selection and screened using primers designed upstream and downstream to the cut site, PCR amplified and sequenced. Clones were further confirmed for knockout of PBRM1 using western blotting.

### Whole cell lysis and Sub-Cellular Fractionation

To make whole cell lysates, cells (70–80% confluent) were washed and collected by scraping into ice-cold PBS, pelleted by centrifugation at 4°C, resuspended in 1X lysis buffer (20mM Tris, pH, 7.5, 150mM NaCl, 1mM EDTA, 1mM EGTA, 1% Triton-X-100, 2.5mM sodium pyrophosphate and 1mM β-glycerophosphate, 1mM Na_3_VO_4_ and 1X complete protease inhibitor cocktail (Roche)), sonicated for 5 cycles with 30 seconds on and 30 seconds off per cycle and centrifuged at maximum speed for 10 minutes at 4°C. The pellet was removed and supernatants collected as whole cell extract for immunoblotting and immunoprecipitation analysis for soluble proteins. For subcellular fractionation, the pellet was collected after centrifugation as above, and resuspended in a hypotonic buffer (10 mM HEPES, pH 7.2, 10 mM KCl, 1.5 mM MgCl_2_, 0.1 mM EGTA, 20 mM NaF, and 100 μM Na_3_VO_4_), and disrupted using a manual homogenizer for 15-20 times. Disruption of the cells was confirmed using a hemacytometer. Following homogenization, samples were centrifuged at 1000 rpm, the supernatant collected, as ‘cytoplasmic fraction’ and the pellet as crude nuclei. The crude nuclei were washed with buffer (10 mM Tris-HCl, pH 7.4, 0.1% NP-40, 0.05% sodium deoxycholate, 10 mM NaCl, and 3 mM MgCl_2_) and lysed in high-salt lysis buffer (20 mM HEPES, pH 7.4, 0.5 M NaCl, 0.5% NP-40, and 1.5 mM MgCl_2_) by sonicating for 5 cycles with 30 seconds on/30 seconds off using Biorupter. After centrifugation at 13,000 rpm in 4°C for 5 min, the pellet was discarded, and supernatant was collected as ‘nuclear fraction’. All the lysis and wash buffers contained 1X Complete protease inhibitor cocktail (Roche). The whole cell extract lysates and the nuclear and cytoplasmic fractions were subjected to BCA Protein Assay (Pierce Chemical Co.) to quantify and normalize the protein levels and then subjected to SDS-PAGE followed by immunoblot analysis.

### Peptide Pull down assay

The K40 α-tubulin peptides were custom synthesize from Thermofisher (K40: IQPDGQMPSDKTIGGGDDSFT; K40ac: IQPDGQMPSD [Kac] TIGGGDDSFT; K40me3: IQPDGQMPSD[Kme3] TIGGGDDSFT) with a biotin label at C-terminal. The peptide pull down assays were performed as described previously by Porter et al., with some modification^25^. The Strepatvidin agarose Resins (ThermoFisher) 20 μl was washed three times with Binding Buffer (0.5 mM DTT, 150 mM NaCl, 50 mM Tris), resuspended in 300 μl of Binding Buffer with 2 μg of biotin-labeled tubulin peptide (K40 only, K40-Ac and K40-me3) (ThermoFisher), and samples were rotated at 4 °C for 2 h. Cells were harvested and lysed in IP Buffer (25 mM Tris, pH 8, 300 mM NaCl, 1% Nonidet P-40, 1 mM EDTA, plus protease inhibitors) and sonicated for 5 cycles using 30 seconds on/30 seconds off. The samples were then spun down at 10,000xg for 10 min, 300 μl of lysate was added to the peptide and resin solution and rotated overnight. The samples were washed for 10 min three times in IP Buffer, the resin was resuspended in 1X Sample Buffer and boiled for 7 min at 95°C, and the samples run on 4-15% SDS PAGE gel (Bio-Rad) and processed for immunoblotting analysis.

### Immunoprecipitation and Immunoblotting Analysis

For immunoprecipitation whole cell lysates or fractionated cytoplasmic lysates were incubated with indicated antibodies and Magnetic A/G beads (Thermo Scientific) or GFP-trap overnight at 4°C. The beads were pelleted and washed with cell lysis buffer three times and were heated in 1X denaturing loading buffer for 10 min at 95°C before being loaded into SDS-PAGE. The cell lysates were separated on a 4-15% gel (Bio-Rad), transferred to PVDF membranes and probed with the indicated antibodies. Densitometric analysis for quantification of expression levels was performed using ImageQuantTL software and data were normalized with α-tubulin.

### Microtubule co-sedimentation assay

MT co-sedimentation assay was performed as described previously by Miller et al., with some modifications^26^. HEK293T wild-type and PBRM1 CRISPR KO cells were harvested and rinsed in PBS twice. Cells were resuspended in BRB80 (80 mM PIPES, 1 mM EGTA, 1 mM MgCl2 pH 6.8) supplemented with protease inhibitor cocktail (Complete mini EDTA free, Roche) and 1 mM DTT. After sonication for 40 sec, lysate was centrifuged at 120,000 × g at 4ºC for an hour. To assemble microtubules (MT), cleared lysate was incubated at 37ºC for 25 min in the presence of 10 μM taxol and 1 mM GTP. MT-containing lysates were layered on prewarmed MT-cushion (5% sucrose in BRB80 supplemented with 10 μM taxol and 1 mM GTP) and centrifuged at 80,000 × g at 37ºC for 30 min. Supernatant was collected and mixed with 5x SDS-PAGE sample buffer. MT pellet was resuspended in 1x SDS-PAGE sample buffer. Lysate containing approximately 36 μg of proteins was used as input and the corresponding amount of supernatant and pellet fractions were resolved on an 8% polyacrylamide gel. Proteins were transferred onto a nitrocellulose membrane and stained with ponceau-S prior to western blotting analysis. The primary antibodies used included anti-PBRM1 rabbit polyclonal antibody, anti-p150Glued mouse monoclonal antibody and anti-MAP4 rabbit polyclonal antibody.

### Immunofluorescence staining and Imaging

Cells seeded on coverslips were fixed either by immersion in cold methanol at −20°C for 5 min followed by rehydration in PBS for 5 min or by fixing with 4% paraformaldehyde (PFA) in PBS at room temperature for 15 min followed by permeabilization with 0.5% Triton X-100 in PBS (PBST) for 20 min. Fixed cells were then blocked with 3.75 % Bovine Serum Albumin (BSA) in PBS for 1 hour and incubated in primary antibodies diluted in 3.75 % BSA for overnight at 4°C. Primary antibodies used for immunofluorescence analysis included rabbit anti-PBRM1, Rabbit anti-SMARCA4 antibody, mouse anti-α-tubulin antibody, Rabbit anti-α tubulin, rabbit anti-ARID2 antibody. Following three washes of 10 mins each using 1X PBS, cells were incubated with corresponding Alexa Fluor-labelled secondary antibodies (Invitrogen, Eugene, OR) at a dilution of 1:2000 at RT for an hour. Cells were then washed three times with 1XPBS, with 10 mins incubation in each wash and then post-fixed for 10 mins using 4% PFA to stabilize the signal. Cells were then counterstained with DAPI (Invitrogen, #, diluted 1:4000) for 10 mins to visualize the DNA. Coverslips were mounted in SlowFade Gold Antifade Mountant (Invitrogen, #S36937). Fixed cells were imaged with a CFI Plan Apochromat Lambda 60X oil, 1.4 NA objective and DS-Qi2 camera and mounted on Nikon Eclipse Ti2-E inverted research microscope (Nikon Instruments, Melville, NY) equipped for standard phase contrast and epifluorescence microscopy, as well as for deconvolution. Image acquisition was carried using an Andor Zyla 4.2+ sCMOS high-sensitivity monochrome camera and was driven by Nikon NIS-Elements Advanced Research (AR) image acquisition and analysis software. Acquired images were processed for advanced 3D and 2D deconvolution modules for improved image quality. Eight-bit images were exported, and figures were prepared using Photoshop version CC software (Adobe Systems, Mountain View, CA). To determine the colocalization and intensity of signal a line was drawn between two spindle pole using deconvolustion microscope NIS elements software. Graphical representations and statistical analyses were performed using GraphPad Prism software version 8. Cell phenotypes were scored visually by counting non-overlapping fields in a raster pattern across the coverslip using the Eclipse Nikon Ti2-E inverted research microscope.

For live cell imaging, cells were plated on 35mm glass-bottomed microwell petri dish (MatTek corporation, Ashland, MA) at a confluence of 5X10^5^ and co-reverse transfected with 2μg of GFP-PBRM1 and pBT097-pAAV Ef1a-H2BmRuby2 using Lipofectamine 2000 (Life technologies) for 48hrs in a 37°C incubator supplied with 5% CO2. Then before imaging, media in the dish was changed to Phenol free complete DMEM media supplemented with 10% FBS to ensure no fluorescence interference from the Phenol. The culture dish was then placed into temperature -controlled stage (Tokai Hit, Shizuoka-ken, Japan) prewarmed to 37°C and supplied with 5% CO2. Z-stacks were acquired every one-two minutes using a CFI Plan Apochromat Lambda 100X oil, NA=1.45, mounted on a Nikon Eclipse Ti2-E with Yokogawa W1 Spinning Disk Confocal Microscope and Photometrics Prime 95B sCMOS camera with NIS-Elements AR software. EDF (Extended Depth of Focus) projections of the images were then generated. Measurement of fluorescence intensity was carried out using Nikon’s NIS Element software.

### Statistical analysis

All data values are represented as standard error of the mean (S.E.M). Graph Pad Prism 7.0 software was used for data analysis. Statistical significance was determined by either Student’s t-test or by one-way ANOVA. Unless otherwise noted, every experiment was done with at least three biologically independent replicates. p-value of less than 0.05 were considered statistically significant. Representative western blots and microscopy images were shown from at least three biologically independent replicates that showed similar results.

**Supplementary Video:** Live cell spinning disc confocal microscopy imaging of *PBRM1 CRISPR-KO HEK293T* cells ectopically expressing GFP-PBRM1 and H2B-mRuby2 to visualize the localization of PBRM1 in the mitotic spindle. Representative image (n=3).

## References

1 Hodges, C., Kirkland, J. G. & Crabtree, G. R. The Many Roles of BAF (mSWI/SNF) and PBAF Complexes in Cancer. Cold Spring Harb Perspect Med 6, doi:10.1101/cshperspect.a026930 (2016).

2 Chambers, A. L. et al. The two different isoforms of the RSC chromatin remodeling complex play distinct roles in DNA damage responses. PLoS One 7, e32016, doi:10.1371/journal.pone.0032016 (2012).

3 Mashtalir, N. et al. Modular Organization and Assembly of SWI/SNF Family Chromatin Remodeling Complexes. Cell 175, 1272–1288 e1220, doi:10.1016/j.cell.2018.09.032 (2018).

4 Kadoch, C. & Crabtree, G. R. Mammalian SWI/SNF chromatin remodeling complexes and cancer: Mechanistic insights gained from human genomics. Sci Adv 1, e1500447, doi:10.1126/sciadv.1500447 (2015).

5 Kadoch, C. et al. Proteomic and bioinformatic analysis of mammalian SWI/SNF complexes identifies extensive roles in human malignancy. Nat Genet 45, 592–601, doi:10.1038/ng.2628 (2013).

6 Gu, Y. F. et al. Modeling Renal Cell Carcinoma in Mice: Bap1 and Pbrm1 Inactivation Drive Tumor Grade. Cancer Discov 7, 900–917, doi:10.1158/2159-8290.CD-17-0292 (2017).

7 Varela, I. et al. Exome sequencing identifies frequent mutation of the SWI/SNF complex gene PBRM1 in renal carcinoma. Nature 469, 539–542, doi:10.1038/nature09639 (2011).

8 Chiang, Y. C. et al. SETD2 Haploinsufficiency for Microtubule Methylation Is an Early Driver of Genomic Instability in Renal Cell Carcinoma. Cancer Res 78, 3135–3146, doi:10.1158/0008-5472.CAN-17-3460 (2018).

9 Park, I. Y. et al. Dual Chromatin and Cytoskeletal Remodeling by SETD2. Cell 166, 950–962, doi:10.1016/j.cell.2016.07.005 (2016).

10 Tang, L., Nogales, E. & Ciferri, C. Structure and function of SWI/SNF chromatin remodeling complexes and mechanistic implications for transcription. Prog Biophys Mol Biol 102, 122–128, doi:10.1016/j.pbiomolbio.2010.05.001 (2010).

11 Baetz, K. K., Krogan, N. J., Emili, A., Greenblatt, J. & Hieter, P. The ctf13-30/CTF13 genomic haploinsufficiency modifier screen identifies the yeast chromatin remodeling complex RSC, which is required for the establishment of sister chromatid cohesion. Mol Cell Biol 24, 1232–1244, doi:10.1128/mcb.24.3.1232-1244.2003 (2004).

12 Huang, J., Hsu, J. M. & Laurent, B. C. The RSC nucleosome-remodeling complex is required for Cohesin’s association with chromosome arms. Mol Cell 13, 739–750, doi:10.1016/s1097-2765(04)00103-0 (2004).

13 Xue, Y. et al. The human SWI/SNF-B chromatin-remodeling complex is related to yeast rsc and localizes at kinetochores of mitotic chromosomes. Proc Natl Acad Sci U S A 97, 13015–13020, doi:10.1073/pnas.240208597 (2000).

14 Brownlee, P. M., Chambers, A. L., Cloney, R., Bianchi, A. & Downs, J. A. BAF180 promotes cohesion and prevents genome instability and aneuploidy. Cell Rep 6, 973–981, doi:10.1016/j.celrep.2014.02.012 (2014).

15 Even, A. et al. ATAT1-enriched vesicles promote microtubule acetylation via axonal transport. Sci Adv 5, eaax2705, doi:10.1126/sciadv.aax2705 (2019).

16 Schaedel, L. et al. Lattice defects induce microtubule self-renewal. Nat Phys 15, 830–838, doi:10.1038/s41567-019-0542-4 (2019).

17 Owen-Hughes, T., Utley, R. T., Cote, J., Peterson, C. L. & Workman, J. L. Persistent site-specific remodeling of a nucleosome array by transient action of the SWI/SNF complex. Science 273, 513–516, doi:10.1126/science.273.5274.513 (1996).

18 Schwabish, M. A. & Struhl, K. The Swi/Snf complex is important for histone eviction during transcriptional activation and RNA polymerase II elongation in vivo. Mol Cell Biol 27, 6987–6995, doi:10.1128/MCB.00717-07 (2007).

19 Sinha, M., Watanabe, S., Johnson, A., Moazed, D. & Peterson, C. L. Recombinational repair within heterochromatin requires ATP-dependent chromatin remodeling. Cell 138, 1109–1121, doi:10.1016/j.cell.2009.07.013 (2009).

20 Larkin, J., Goh, X. Y., Vetter, M., Pickering, L. & Swanton, C. Epigenetic regulation in RCC: opportunities for therapeutic intervention? Nat Rev Urol 9, 147–155, doi:10.1038/nrurol.2011.236 (2012).

21 Turajlic, S. et al. Tracking Cancer Evolution Reveals Constrained Routes to Metastases: TRACERx Renal. Cell 173, 581–594 e512, doi:10.1016/j.cell.2018.03.057 (2018).

22 Turajlic, S. et al. Deterministic Evolutionary Trajectories Influence Primary Tumor Growth: TRACERx Renal. Cell 173, 595–610 e511, doi:10.1016/j.cell.2018.03.043 (2018).

23 Gong, F. et al. Screen identifies bromodomain protein ZMYND8 in chromatin recognition of transcription-associated DNA damage that promotes homologous recombination. Genes Dev 29, 197–211, doi:10.1101/gad.252189.114 (2015).

24 Joo, E. E. & Yamada, K. M. MYPT1 regulates contractility and microtubule acetylation to modulate integrin adhesions and matrix assembly. Nat Commun 5, 3510, doi:10.1038/ncomms4510 (2014).

25 Porter, E. G. & Dykhuizen, E. C. Individual Bromodomains of Polybromo-1 Contribute to Chromatin Association and Tumor Suppression in Clear Cell Renal Carcinoma. J Biol Chem 292, 2601–2610, doi:10.1074/jbc.M116.746875 (2017).

26 Miller, L. M. et al. Methods in tubulin proteomics. Methods Cell Biol 95, 105–126, doi:10.1016/S0091-679X(10)95007-3 (2010).

